# Fully Automated Detection of Paramagnetic Rims in Multiple Sclerosis Lesions on 3T Susceptibility-Based MR Imaging

**DOI:** 10.1101/2020.08.31.276238

**Authors:** Carolyn Lou, Pascal Sati, Martina Absinta, Kelly Clark, Jordan D. Dworkin, Alessandra M. Valcarcel, Matthew K. Schindler, Daniel S. Reich, Elizabeth M. Sweeney, Russell T. Shinohara

## Abstract

**Background and Purpose:** The presence of a paramagnetic rim around a white matter lesion has recently been shown to be a hallmark of a particular pathological type of multiple sclerosis (MS) lesion. Increased prevalence of these paramagnetic rim lesions (PRLs) is associated with a more severe disease course in MS. The identification of these lesions is time-consuming to perform manually. We present a method to automatically detect PRLs on 3T T2*-phase images.

**Methods:** T1-weighted, T2-FLAIR, and T2*-phase MRI of the brain were collected at 3T for 19 subjects with MS. The images were then processed with lesion segmentation, lesion center detection, lesion labelling, and lesion-level radiomic feature extraction. A total of 877 lesions were identified, 118 (13%) of which contained a paramagnetic rim. We divided our data into a training set (15 patients, 673 lesions) and a testing set (4 patients, 204 lesions). We fit a random forest classification model on the training set and assessed our ability to classify lesions as PRL on the test set.

**Results:** The number of PRLs per subject identified via our automated lesion labelling method was highly correlated with the gold standard count of PRLs per subject, r = 0.91 (95% CI [0.79, 0.97]). The classification algorithm using radiomic features can classify a lesion as PRL or not with an area under the curve of 0.80 (95% CI [0.67, 0.86]).

**Conclusion:** This study develops a fully automated technique for the detection of paramagnetic rim lesions using standard T1 and FLAIR sequences and a T2*phase sequence obtained on 3T MR images.

**Highlights:**

- A fully automated method for both the identification and classification of paramagnetic rim lesions is proposed.
- Radiomic features in conjunction with machine learning algorithms can accurately classify paramagnetic rim lesions.
- Challenges for classification are largely driven by heterogeneity between lesions, including equivocal rim signatures and lesion location.

## Introduction

Multiple sclerosis (MS) is a demyelinating and inflammatory disease of the central nervous system whose hallmark is lesions in the brain and spinal cord (1). These lesions can be detected *in vivo* with magnetic resonance imaging (MRI) and are often quantified as total lesion volume and lesion count, both of which can be used as measures of disease burden and to track disease progression (2). Imaging biomarkers such as these are commonly used in the clinic and as surrogate endpoints in clinical trials (3,4). However, other known biological processes of MS are left uncaptured.

Chronic active lesions, which are a subset of MS lesions that are more prevalent in patients with more severe disease (5–7), have imaging and histopathology findings suggestive of ongoing tissue damage (8–10) and have until recently only been detectable by histopathology. These lesions have been variously termed chronic active, slowly expanding, or smoldering lesions. At an estimated prevalence of 10-15% of all MS lesions, this type of lesion is sufficiently common and deleterious to warrant considerable efforts for biomarker development (6,8,9,11). On T2*-phase MRI contrast, they are identifiable by curvilinear hypointensity along the edge of the lesion that corresponds with of iron laden phagocytic cells observed on histopathological specimens (8,9,12). These lesions have been variously termed chronic active, slowly expanding, or smoldering lesions. Here, we refer to these lesions as paramagnetic rim lesions (PRLs).

When first observed on MRI, the rim of a PRL was only visible on scans from ultra-high-field strength (7T) magnets (13–16). Recently, PRLs have been shown to be identifiable on the more commonly available high-field strength (3T) MRI scans as well, albeit with lower inter- and intra-rater reliability (17). This development strengthens their viability as a target on clinical MRI protocols, particularly because the sequences studied can be acquired with high spatial resolution in less than 4 minutes (18). Previous studies of the PRLs have noted the geometric nature of the rim and worked to identify the rim on the quantitative susceptibility mapping (QSM) contrast as well (19–21).

Because visually inspecting every MS lesion for the presence of a paramagnetic rim is difficult, time consuming, and prone to inter- and intra-rater variability, an automated method for identifying PRLs would improve efficiency and facilitate translation of this imaging biomarker into larger research studies and clinical practice. One way to identify PRLs is through quantification of visual patterns that objectively characterize these data, which can be accomplished through radiomics. Radiomics is an emerging field of research that encompasses the extraction of quantitative features from biomedical images that may reflect underlying pathophysiology (22). It has been shown to be a useful tool in the analysis of chest CT scans (23,24) and MR images (25,26). Studies have shown that radiomic features are often useful predictors of, or are associated with, known hallmarks of disease, although they have not been used extensively in the MS literature. Here, we use radiomic features along with a random forest classification model, which can flexibly model high dimensional data. Our method is fully automated and uses a T2*-phase volume with isometric voxels and high spatial resolution that is acquired in a clinically feasible acquisition time at 3T (18).

## Materials and Methods

### Study population

We studied 19 subjects with MS who were scanned under an institutional review board– approved natural history protocol at the National Institutes of Health (NIH). Subjects’ age at the time of scanning ranged from 20 to 66 years, with a mean age of 45 years (sd = 12) (Table 1).

**Table 1:** Demographics of Study Sample

| Demographics |  |
| --- | --- |
| N | 19 |
| Age (mean (SD)) | 45 (12) |
| Male (%) | 8 (42) |
| Phenotype (%) |  |
| Primary Progressive MS | 3 (16) |
| Relapsing-Remitting MS | 11 (58) |
| Secondary Progressive MS | 5 (26) |
| Disease duration in years (mean (SD)) | 14.6 (9.1) |
| EDSS (median (range)) | 2.5 (1.0—7.0) |
| Treatment |  |
| untreated | 5 (26) |
| glatiramer acetate | 1 (5) |
| interferon beta-1a | 4 (21) |
| dimethyl fumarate | 6 (32) |
| fingolimod | 1 (5) |
| natalizumab | 1 (5) |
| rituximab | 1 (5) |

Written informed consent was obtained from all participants. Data from this study can be shared upon reasonable request and completion of a Data Transfer Agreement with the National Institutes of Health.

### MR Imaging acquisition

All subjects were imaged on a Siemens Magnetom Skyra (Siemens, Erlangen, Germany) 3T scanner, using a body transmit coil and a 32-channel receive array coil, at the National Institutes of Health in Bethesda, Maryland. Imaging acquisition included the following sequences:

- a whole-brain 3D T2-weighted fluid-attenuated inversion recovery (FLAIR) sequence (repetition time, TR = 4800 ms; echo time, TE = 354 ms; inversion time, TI = 1800 ms; flip angle, FA = 120°; acquisition time, TA = 6 minutes 30 seconds; 256 axial slices; 1mm isometric voxel resolution);
- a whole-brain 3D T1-weighted magnetization-prepared rapid gradient echo (T1) sequence (TR = 7.8 ms; TE = 3 ms; FA = 18°; TA = 3 minutes 35 seconds; 256 sagittal slices; 1mm isometric voxel resolution), and
- a 3D segmented echo-planar imaging (EPI) sequence with whole-brain coverage providing T2* magnitude and phase contrasts (TR = 64 ms; TE = 35 ms; flip angle, FA = 10°; TA = 5 minutes 46 seconds; 251 sagittal slices; 0.65mm isometric voxel resolution).

Additional standard MRI sequences, including a postcontrast 3D T1-weighted MPRAGE sequence for the identification of gadolinium-enhancing lesions, were also acquired.

### Manual paramagnetic rim lesion assessment

Supratentorial non-gadolinium enhancing MS lesions were visually inspected for the presence of a paramagnetic rim on T2* magnitude and unwrapped phase images by a neurologist with 14 years of experience in neuroimaging science (5,13,17). As previously described (27), a PRL is identified when a hypointense signal on phase images is observed surrounding the periphery of the lesion, while being either hyper- or isointense in its inner portion.

### Image preprocessing

Phase images were unwrapped and filtered as previously described (13). T1, FLAIR, and phase images were then preprocessed using the *fslr* R package (28), an R wrapper for the FSL software (29,30). Images were visualized with ITK-SNAP (31). The T2* magnitude contrast was not used in this method.

We first applied the N4 inhomogeneity correction algorithm (32). We then rigidly registered both the T1 and the FLAIR images to the T2*-phase image space, resampling to 0.65 mm isometric resolution and using a mutual information cost function and nearest neighbor interpolation. We used multi-atlas skull stripping (MASS) to identify cerebral tissue in the images in T1 space (33). In two cases, MASS yielded poorly skull-stripped images based on visual inspection. For those two cases, we instead used the FSL brain extraction tool for skull-stripping (29). As a final step, we performed WhiteStripe intensity normalization on the otherwise preprocessed T1, FLAIR, and phase images (34).

### Lesion labelling

Our lesion labelling method relies on access to maps that represent voxel-wise probabilities of being a lesion, so we chose the automatic lesion segmentation method MIMoSA both for its ability to integrate multimodal information and for its ability to provide voxel-level probability maps (35). Manual lesion segmentation was conducted by a research assistant with 1 year of experience, who was trained by a board-certified neurologist with extensive expertise in neuroimmunology and MRI.

We trained the MIMoSA algorithm (36) with the manual segmentations as a gold standard and the T1 and FLAIR images as input. We implemented a leave-one-out cross-validation approach, where data from all but one subject was used to train a MIMoSA model, and that model was subsequently applied to the remaining subject. This was done for every subject in our cohort. When parallelized across 8 cores of a CPU of an Intel(R) Xeon(R) E5-2699 v4 @ 2.20GHz processor in a high-performance computing environment, training a single model on 19 subjects took approximately 6 hours.

From each k-fold model, we extracted probability maps that contain voxel-wise probabilities of being a white matter lesion. We then binarized these probability maps into lesion segmentation maps via a subject-specific estimated optimal threshold that was identified out of a user-provided range of possible thresholds and then chosen based on amount of overlap with a gold-standard lesion segmentation as measured by a Sørensen-Dice coefficient (37). Because our lesion segmentation masks did not always cover the entire area of a lesion, we then dilated the masks by one voxel in each direction to increase the likelihood of detecting the paramagnetic rim signal, which occurs on the boundary of lesions.

After lesion segmentation masks were obtained, we used the lesion probability maps as input to a center detection method (38) to identify distinct lesions based on the texture of the lesion tissue. We then used a nearest-neighbor approach to classify the remainder of the lesion segmentation map into those identified lesions (Figure 1). At this point, we assigned PRL status to the identified lesions based on the presence of any overlap with the manual PRL labels described previously.

**Figure 1.**
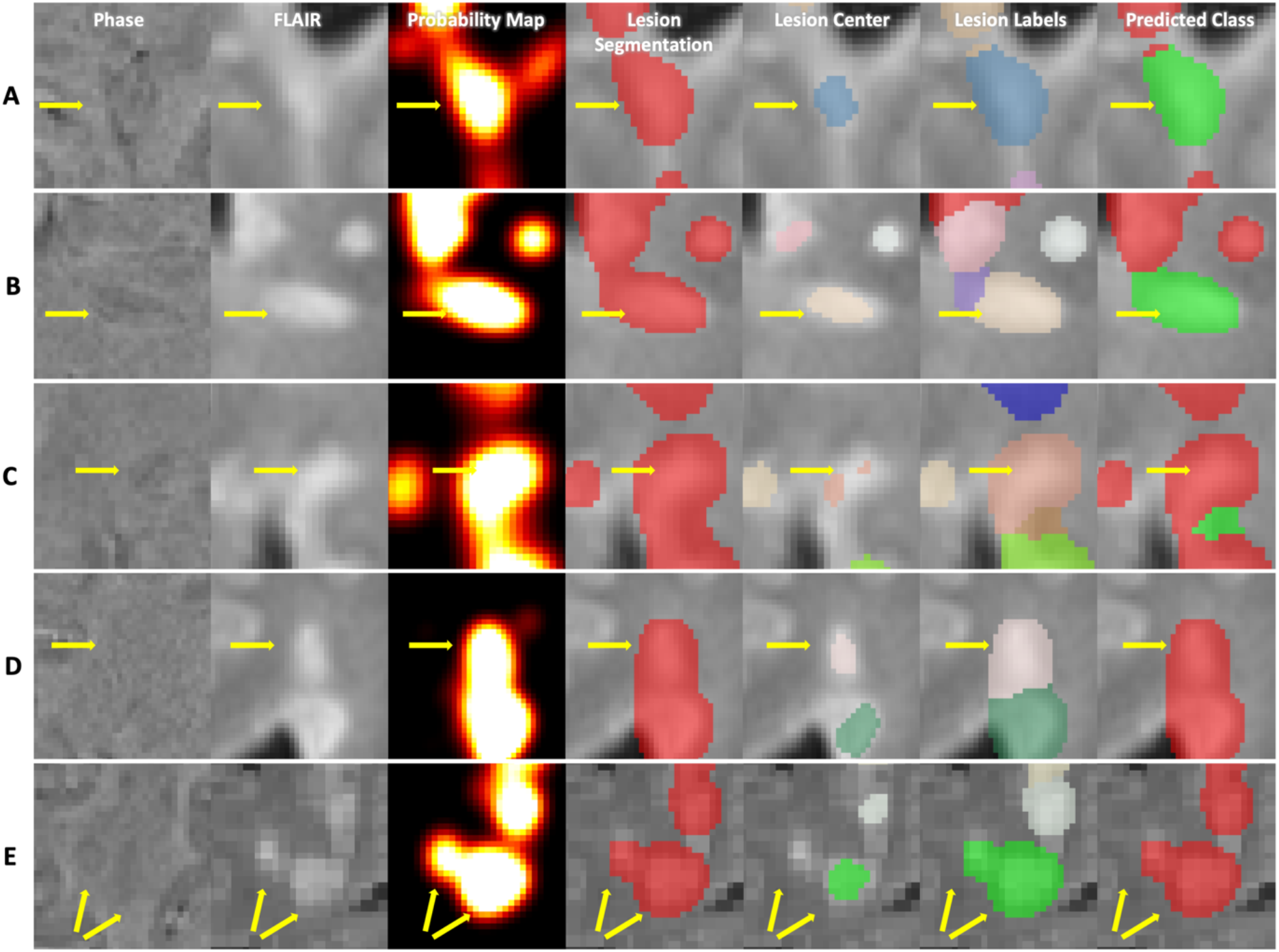
A visualization of the steps of the method for five different lesions. Each column corresponds to a different part of the method, and each row corresponds to a different lesion of interest. In the last column, lesions classified as PRLs are visualized as green, and lesions classified as not PRLs are visualized as red. Subfigure A shows a lesion that was both identified as a PRL and classified as a PRL, i.e. a true positive. Subfigure B shows a lesion that was identified as not a PRL but classified as a PRL, i.e. a false positive. Correspondingly, subfigure C shows a false negative lesion, and subfigure D shows a true negative lesion. Subfigure E shows a lesion that was automatically labelled as a single lesion but is actually a confluence of lesions.

Due to failures in the lesion labelling process, a subset of abnormalities automatically identified by our method might, to a manual rater, be considered clusters of confluent lesions. Because we did not have access to manual segmentations of distinct lesions, we instead relied on a combination of our lesion labelling method and connected components analysis to label lesions as confluent. Specifically, if connected components only identified one cluster where our lesion labelling method identified more than one lesion, we labelled the constituent lesions as confluent.

### Radiomics image analysis

For lesions that were identified with our automatic pipeline, we conducted a radiomics analysis to characterize each lesion with intensity-based statistics only on the phase contrast (39,40). These include 44 features that summarize the intensities in an individual lesion with measures that can be described in 3 general ways: statistics that describe the average and spread of the intensities, statistics that describe the shape of the distribution of intensities, and statistics that describe the diversity of intensities (40). For example, features like the mean, defined as 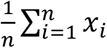, and interquartile range, defined as *abs*(*x*_75%_ – *x*_25%_), are included in the first group, where *x_i_* represents intensity value at voxel *i*. Features like variance, defined as 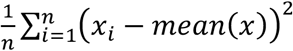, and skew, defined as 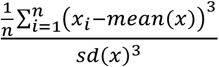, are included in the second group, and features like energy, defined as 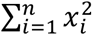, uniformity, defined as 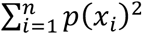, and entropy, defined as 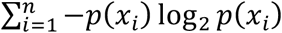, are included in the third group. A full list and detailed equations for each of the first-order radiomic features can be found in the supplemental material of (40).

### Prediction model

The radiomic features were used as candidate predictors in our subsequent prediction modelling for classification of lesions as either being PRL or not. Class labels for each lesion were previously assigned during the lesion labelling step. We split our dataset into a training set and test set by subject, randomly assigning lesions from 15 subjects into the training set and assigning lesions from the remaining 4 subjects into the test set, approximating an 80/20 split. Both sets were examined to ensure that at least 100 lesions were present in each group.

Because PRLs were of a minority class (approximately 13% of the lesions were classified as being a PRL), we used Synthetic Minority Oversampling TEchnique (SMOTE) to synthetically balance our data (41). With SMOTE, we oversampled the minority class, the PRLs, by the reciprocal of the percentage of PRLs present in the dataset, and we did not undersample the majority class. We then trained a random forest classifier, chosen for its ability to flexibly model a large number of features, with 10-fold cross-validation using the R package *caret* (42,43). We summarized performance results using 0.5 as a threshold where applicable. We also derived empirical confidence intervals for those measurements by randomly reassigning the training and test set and repeating the above process 1000 times. We assessed variable importance in the random forest as the percent increase in mean-squared error for a model with the variable over a model with a permuted version of that variable. We then scaled that measure for comparability across variables.

### Post-hoc analysis

An additional board-certified neurologist (MS) with extensive expertise in neuroimmunology and MRI, who was not involved in the generation of the manual PRL labels, examined each misclassified lesion. We rated lesions on a 5-point scale, where 1 indicated definitely not a PRL, 2 indicated probably not a PRL, 3 indicated uncertain, 4 indicated probably a PRL, and 5 indicated definitely a PRL. Some lesions were automatically labelled as one lesion but were actually a confluence of lesions (Figure 1). We assigned manual ratings to these confluent clusters based on the presence of at least one PRL. We also assessed the method’s performance only for lesions that were not part of a confluent cluster.

## Results

The final dataset included a total of 877 lesions in 19 subjects identified by our automated lesion labelling method, 118 (13%) of which we found to be PRLs by overlap with the manual annotation. The average number of lesions per subject was 46.2 (sd = 19.8), and the average number of PRLs per subject was 6.2 (sd = 4.0). Table 2 summarizes by subject the total number of lesions identified from our lesion labelling method, the number of PRLs identified from our lesion labelling method, and the number of PRLs identified by a manual rater. The number of identified PRLs by our method was highly correlated with the gold standard count of PRLs, r = 0.91 (95% CI [0.79, 0.97]) (Figure 2).

**Figure 2.**
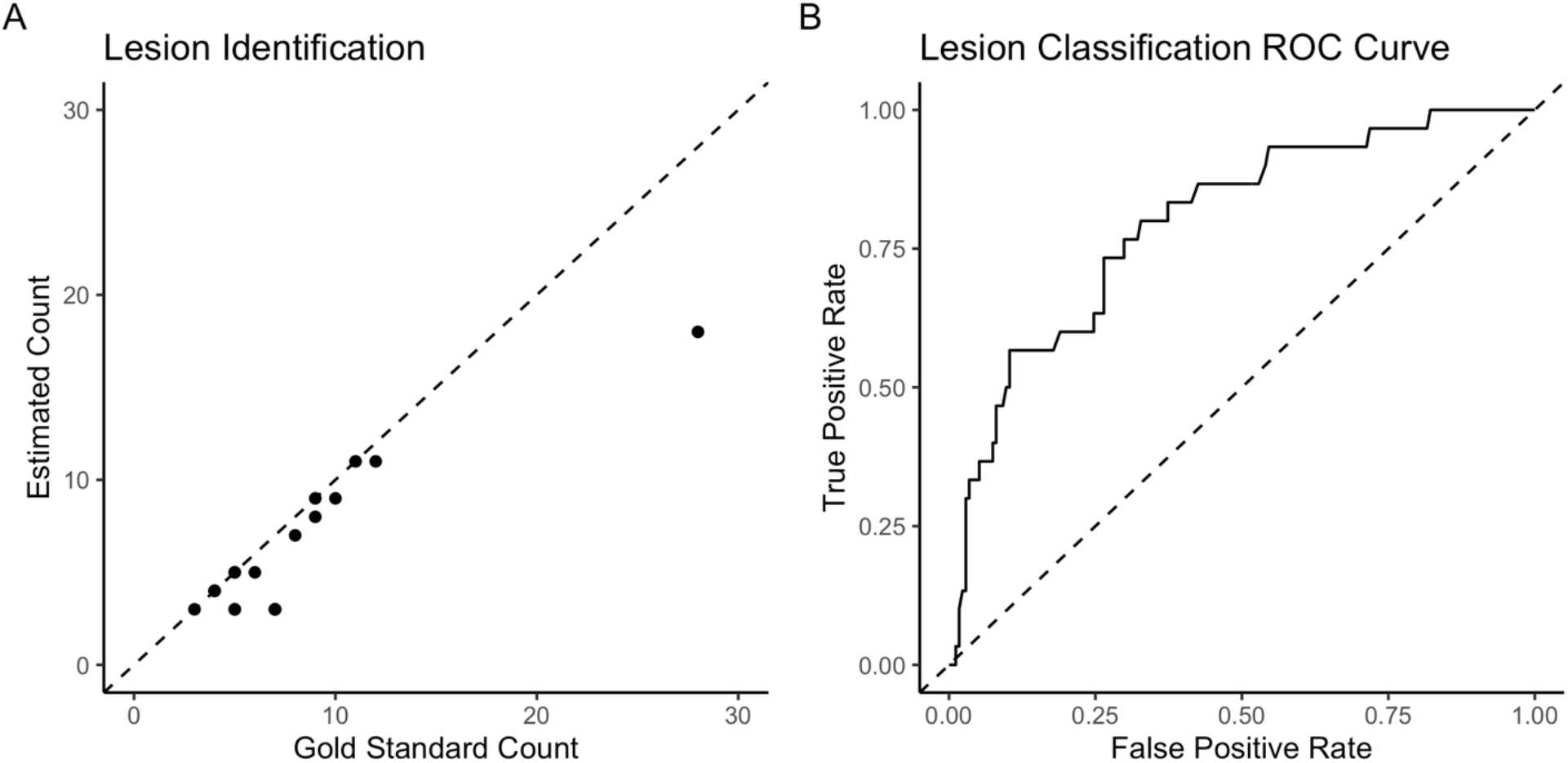
Subfigure A shows the gold standard count of PRLs against the number of PRLs identified via our lesion identification method, r = 0.91 (0.79, 0.97). Subfigure B shows the ROC curve after classification, AUC = 0.80 (0.67, 0.86).

**Table 2:**
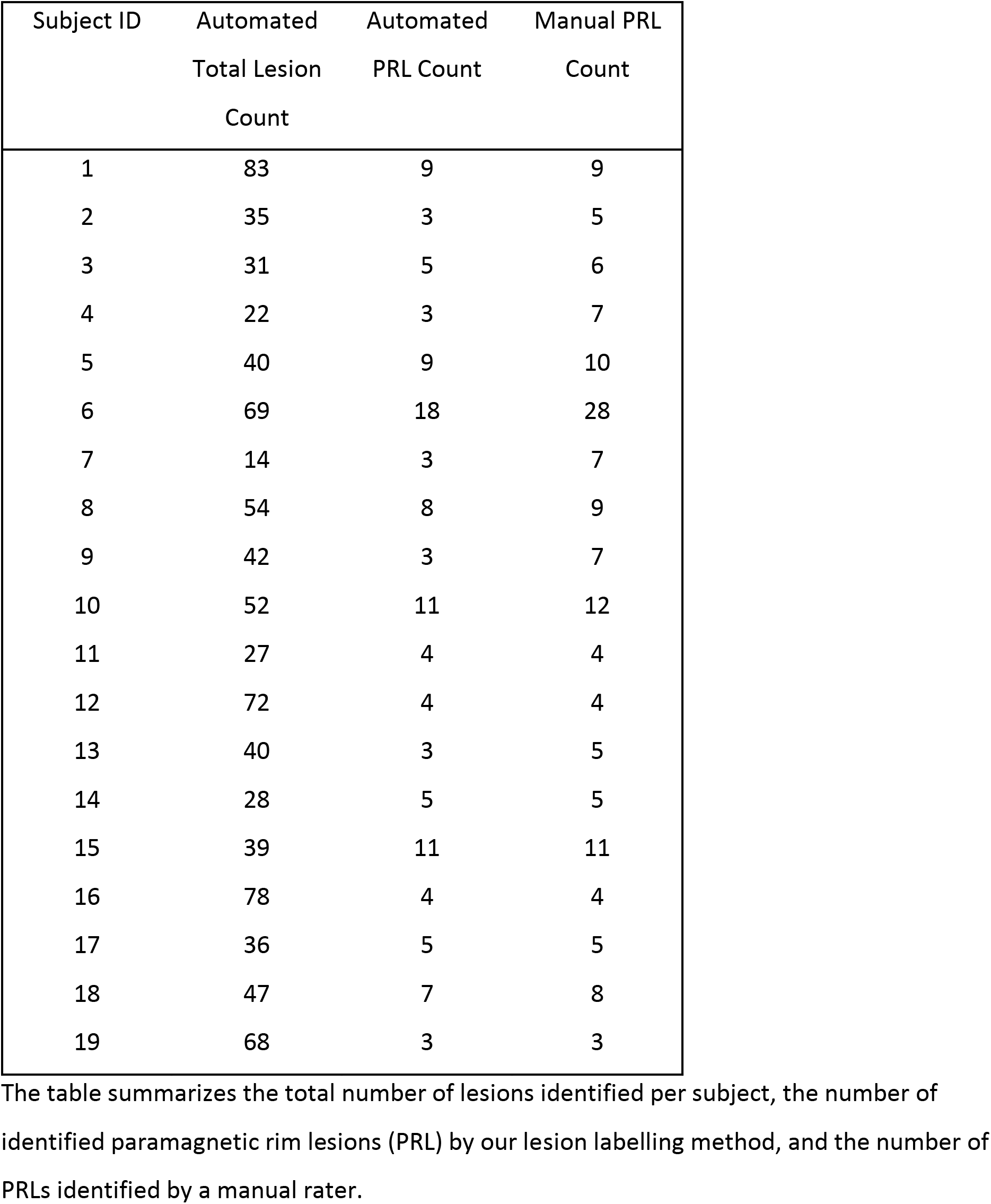
Lesion Counts by Subject

The table summarizes the total number of lesions identified per subject, the number of identified paramagnetic rim lesions (PRL) by our lesion labelling method, and the number of PRLs identified by a manual rater.

We trained a random forest classification model using PRL status from the lesion labelling method as the label. In the iteration that we used to derive performance measures, there were 673 lesions in the training set, 88 of which were PRLs, and 204 lesions in the testing set, 30 of which were PRLs. We were able to classify lesions as PRL or not with an AUC of 0.80 (95% CI [0.67, 0.86]). Using 0.5 as a probability threshold, 150 lesions were accurately classified as not PRL, 24 lesions were false positives, 13 were false negatives, and 17 were classified correctly as PRL (Table 3). A breakdown of the classification results for the test set lesions by subject is also provided in Table 3, from which we can see that variability in classification accuracy does not seem to be driven by poor performance in a minority of subjects but rather by heterogeneity in the lesions themselves.

**Table 3:** Summary of Classification Performance Measures

| Contingency Table |  |  |  |  |
| --- | --- | --- | --- | --- |
|  | Reference |  |  |  |
| Prediction | Rim Negative |  | Rim Positive |  |
| Rim Negative | 150 |  | 13 |  |
| Rim Positive | 24 |  | 17 |  |
| Performance Measures |  |  |  |  |
| Accuracy |  |  | 0.82 (0.71, 0.86) |  |
| Positive Predictive Value |  |  | 0.41 (0.16, 0.53) |  |
| Negative Predictive Value |  |  | 0.92 (0.87, 0.97) |  |
| False Positive Rate |  |  | 0.14 (0.08, 0.27) |  |
| False Negative Rate |  |  | 0.43 (0.22, 0.72) |  |
| Sensitivity |  |  | 0.57 (0.29, 0.74) |  |
| Specificity |  |  | 0.86 (0.72, 0.92) |  |
| Testing Set Lesion Classification Count by Subject |  |  |  |  |
| Subject | True<br>Negative | False Negative | False<br>Positive | True<br>Positive |
| 1 | 70 | 4 | 4 | 5 |
| 4 | 18 | 2 | 1 | 1 |
| 11 | 30 | 5 | 11 | 6 |
| 20 | 32 | 2 | 8 | 5 |

The table summarizes the performance measures we observed for the classification of lesions as PRLs or not.

We also examined the results of the method for lesions that were not part of a confluent cluster. A total of 62 lesions in the test set were not confluent, and we were able to classify them with an AUC of 0.91. Using 0.5 as a probability threshold, 50 lesions were accurately classified as not PRL, 4 were false positive, 2 were false negative, and 6 were accurately classified as PRL (Table 4). We provide a summary of additional performance measures in Table 4.

**Table 4:**
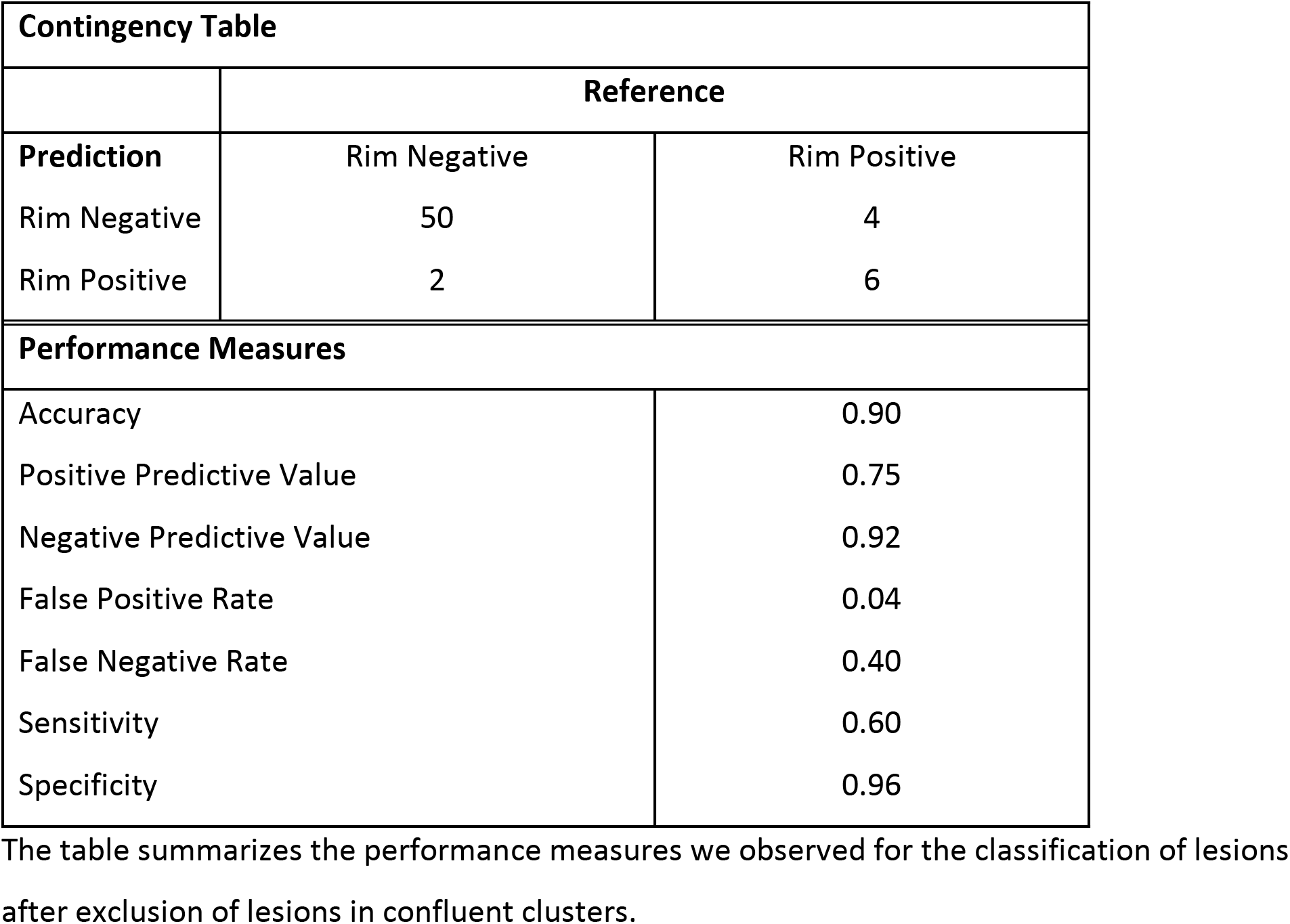
Summary of Classification Performance Measures, Excluding Confluent Lesions

A visualization of lesions that were true positive, false positive, false negative, and true negative respectively is provided in Figure 1. From subfigure B, where we see the method illustrated for a lesion that was identified as a not a PRL but classified as a PRL, we can see that hypointensities can manifest around a lesion even when they cannot be rated as a rim. Conversely, from subfigure C, which shows a lesion that was identified as a PRL but classified as not a PRL, we can see that despite the presence of hypointensities that are visible to the eye, certain PRLs may not display a signal strong enough to be captured by radiomic features.

The random forest identified uniformity, entropy, and energy as the most important radiomic features in classifying lesions, which are all radiomic features that aim to describe the diversity of the data points. (Figure 3). Other radiomic features that were important were mode, kurtosis, skew, geometric mean, and quantile features. Entropy and uniformity were both higher in lesions that were not PRL, and energy was higher in lesions that were PRL.

**Figure 3.**
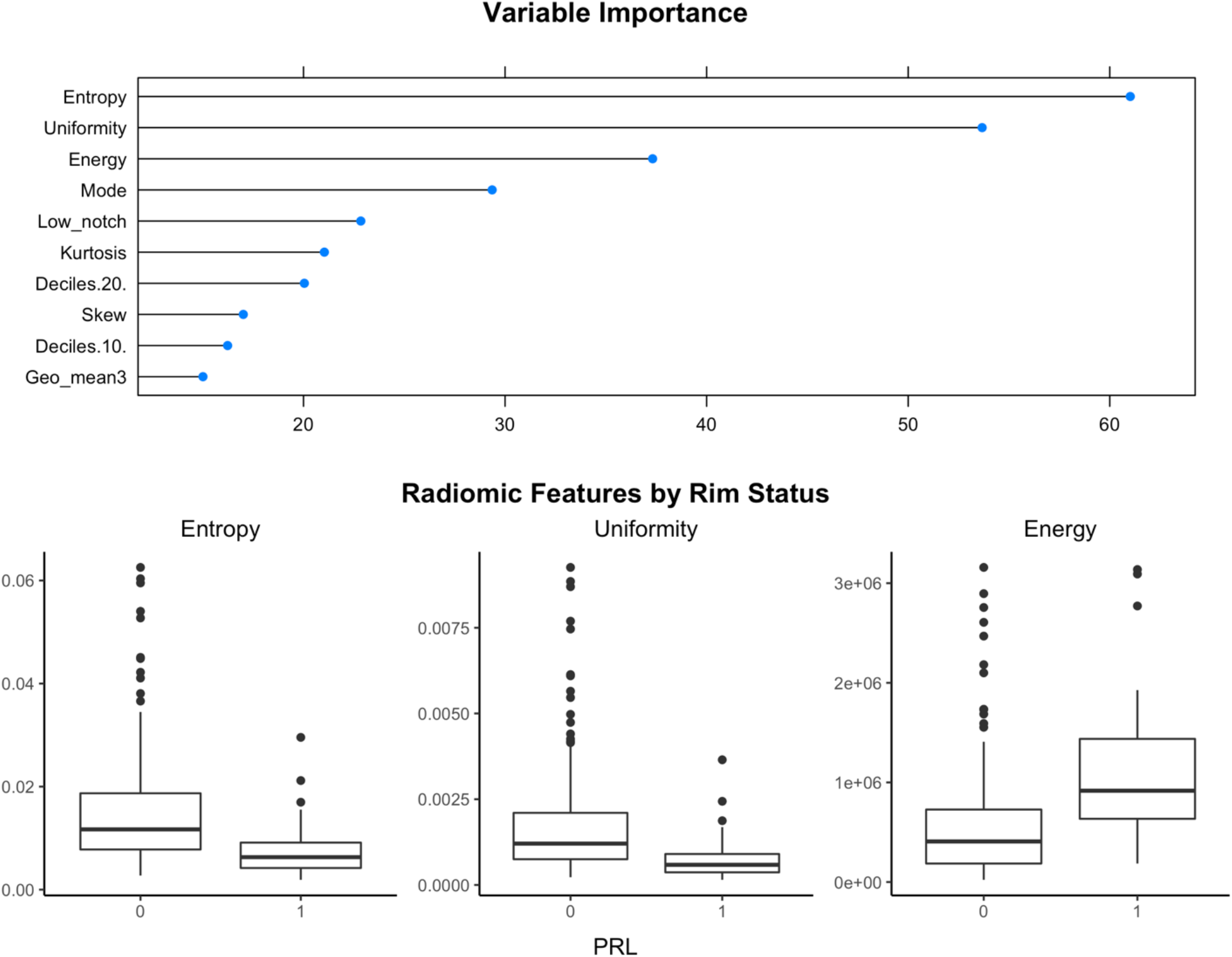
The variables identified as the most important by our model for determining the presence of PRLs were entropy, uniformity, and energy. Here, we measure variable importance as the percent increase in mean squared error for the model with the variable over the model with a permuted version of that variable, scaled for comparability across variables. Boxplots of entropy, uniformity, and energy on the lesions from the test set show that PRLs and non-PRLs seem to differ on those measures, supporting the theory that they are important for distinguishing the two kinds of lesions.

A second expert manually rated the 37 lesions that were misclassified by the model. The rater deemed that 3 lesions included too much artifact to assess PRL status, and 18 lesions were confluent lesions. Of the lesions not part of a confluent cluster, 10 were false positive lesions and 6 were false negative lesions. Of those 10 false positive lesions, 2 of these lesions were rated as definitely a PRL, 2 were rated as probably a PRL, 4 were rated as probably not a PRL, and 2 were rated as definitely not a PRL. For the 6 false negative lesions that were not confluent, 5 were rated as definitely a PRL and 1 was rated as definitely not a PRL.

As for confluent clusters, 11 were deemed false positives and 7 were false negatives. These were rated according to the presence of at least one PRL in each confluent cluster. Of the 11 false positive lesions, 4 were rated as definitely not a PRL, 3 were rated as probably a PRL, and 4 were rated as definitely a PRL. Of the 7 false negative lesions, 2 were rated as probably a PRL and 5 were rated as definitely a PRL. Note that the confluence defined here was as judged by the manual rater. This differs from but complements the confluence definition employed for the primary test set analysis, which was the definition based on the automated analysis used to derive the performance measures reported in Table 4.

## Discussion

Preliminary studies have shown that the existence of a paramagnetic rim around an MS lesion is an important biomarker with possible clinical implications, shown to be indicative of chronic inflammation, associated with heightened disability, and resistant to current disease-modifying treatments (5). However, paramagnetic rims are time-consuming to identify manually, even by highly trained experts (17). In this paper, we developed a fully automatic method for the detection of the paramagnetic rim on a 3T MRI using a submillimeter isometric, clinically feasible, segmented-EPI sequence (17,18). Automation of PRL identification that relies on objective assessment will aid larger scaled studies assessing this promising imaging biomarker in MS.

The proposed method relies on radiomics for automated PRL identification and classification. Radiomic features have been used previously in other contexts, but none were used specifically to classify PRLs. The radiomic features that were the most important in this context aimed to measure the variability of intensity within a lesion (entropy and uniformity) or quantify the magnitudes of the intensities themselves (energy).

Both entropy and uniformity are measures based on the probability of observing a particular intensity within a lesion. Because we did not bin the voxel intensities, this probability of observing a particular intensity is fairly low, which is reflected in the observed range of uniformity in this study. Uniformity is a direct measure of homogeneity of the intensities within a lesion. We expect uniformity to be lower for PRLs due to the presence of both intensities representing normal appearing tissue and hypointensities from the paramagnetic rim. Lesions that are not PRLs do not appear with any distinct signature on the phase image, leading to a higher uniformity.

Entropy takes the probability of observing a particular intensity within a lesion and transforms it such that the measure reflects the amount of variation observed. Because of the aforementioned lack of binning, entropy here more accurately reflects lesion size in that given our more homogenous set of probabilities, a smaller probability of observing a given intensity results in a smaller measure of entropy, and larger lesions yield a smaller probability of observing a given intensity. In this dataset, PRLs tend to have smaller values of entropy, possibly reflecting a larger size.

Energy is a measure of the magnitude of intensities within a lesion. Here, PRLs manifest with higher energy because of the way the phase image was created and the subsequent range of the intensities. Hypointensities on the phase image used in this study represent more extreme negative values instead of values closer to 0, with more extreme hypointensities resulting in more extreme energy values.

Many of the lesions that the model misclassified were confluent lesions that were labelled as a single lesion. While the percentages of confluent lesions among correctly classified lesions was 66%, the percentage of confluent lesions among incorrectly classified lesions was 84%, suggesting that confluence negatively influences the model’s ability to classify lesions as PRL or not. We provide an example of one of these confluent lesions in Figure 1, Subfigure E. In this lesion, although one of the encompassed lesions contained a clear rim signal, the larger of the two does not. Because the majority of the voxels included in the confluent lesion belong to the encompassed one without a rim signal, the first-order radiomic features extracted from this confluent lesion reflected that signal.

Artifact made it difficult for a manual rater to rate some of the lesions; our model typically (perhaps incorrectly) rated these as PRL. Of the “false positive” lesions, as determined by the initial PRL manual delineations, while half of those were separately rated as definitely or probably not a PRL, half were rated as definitely or probably a PRL. We also note that for the false positives, around half of the manual ratings were between 2 and 4 on a 5-point scale indicating that even for an expert rater, a large portion of these lesions were difficult to classify. Of the false negative lesions, almost all were rated as definitely a PRL.

We dilated our lesion segmentation map to increase the likelihood that a rim signal would be included in a lesion label. Because of this artificial augmentation, periventricular lesions and lesions closer to the cortex could be difficult to classify due to inclusion of non-lesional phase-hypointensities in a lesion map, such as ventricles or cortical tissue.

These issues could be addressed by taking a more nuanced approach to modelling the probability of having a rim. Here, we treated the identification of PRLs as a binary classification problem, invoking a random forest to predict if a given lesion was a PRL or not. However, the identification of PRLs can be difficult because of the myriad of factors that drive the clarity and strength of a rim signature, some of which are technical and some of which reflect biological processes. As noted in Figure 1, while some lesions exhibit a rim unequivocally, other lesions exhibit a more equivocal signature. This renders the task of rating lesions as PRL or not difficult, both for manual raters and automated classifiers. In fact, previous research has shown that intra- and interrater reliability for paramagnetic rim evaluation are substantial but not perfect, with a Cohen *κ* of 0.77 and 0.71 respectively (17). A future, more nuanced approach could treat the presence of a rim as a continuous measure instead of a binary classification. This would likely more accurately reflect underlying biological processes as well, as the amount of iron-containing phagocytes at the edge of a lesion can vary across lesions (8).

### Limitations

A major limitation to current assessments of paramagnetic rims is that no international consensus exists on criteria for determining this imaging signature. This limitation may hinder the application of the proposed methodology to new studies in which differing definitions of paramagnetic rims may be desired based on local practices. While signal-to-noise ratio is higher on a 7T MR image, allowing for higher inter- and intra-rater reliability, they remain low across contrast types on 3T (17). However, our study relies on techniques that perform well on 3T images, so extensions to 7T would require additional validation.

In addition, PRLs are a less common type of lesion. In the current study, 13% of the lesions that were identified had rims. Because they are a rare event, classical machine learning models may need to be adjusted in order to classify them with appropriate consideration. In the current study, we employed SMOTE to artificially balance our training data. Other machine learning methods may benefit more from other solutions.

## Conclusion

This study introduces a fully automated method for the identification and classification of paramagnetic rim lesions relying solely on 3T MR images, which are commonly available in a clinical setting. Automation of this process is important for the continued development of the scientific community’s knowledge around these lesions and their implications for disease burden.

## Funding

Dr. Pascal Sati, Dr. Martina Absinta, and Dr. Daniel S. Reich are supported by the Intramural Research Program of the National Institute of Neurological Disorders and Stroke, National Institutes of Health, Bethesda, Maryland, USA. Dr. Martina Absinta is supported by the Conrad N. Hilton Foundation (grant#17313). Dr. Schindler is supported by the National Center for Advancing Translational Sciences of the National Institutes of Health under award number KL2TR001879. Ms. Lou and Dr. Shinohara are supported by awards R01NS112274 and R01NS060910 from the National Institute of Neurological Disorders and Stroke, and R01MH112847 from the National Institute of Mental Health. The content is solely the responsibility of the authors and does not necessarily represent the official views of the National Institutes of Health.

## Declarations of interest

None

## Acknowledgments

We thank the NINDS Neuroimmunology Clinic for recruiting and assessing the patients.

## Notes

### Competing Interest Statement

The authors have declared no competing interest.

